# Temporal dynamics of cholinergic activity in the septo-hippocampal system

**DOI:** 10.1101/2022.06.12.495208

**Authors:** Jeffrey D. Kopsick, Kyle Hartzell, Hallie Lazaro, Pranav Nambiar, Michael E. Hasselmo, Holger Dannenberg

## Abstract

Cholinergic projection neurons in the medial septum and diagonal band of Broca are the major source of cholinergic modulation of hippocampal circuit functions that support neural coding of location and running speed. Changes in cholinergic modulation are known to correlate with changes in brain states, cognitive functions, and behavior. However, whether cholinergic modulation can change fast enough to serve as a potential speed signal in hippocampal and parahippocampal cortices and whether the temporal dynamics in such a signal depend on the presence of visual cues remain unknown. In this study, we use a fiber-photometric approach to quantify the temporal dynamics of cholinergic activity in freely moving mice as a function of the animal’s running speed and visual cues. We show that the population activity of cholinergic neurons in the medial septum and diagonal band of Broca changes fast enough to be aligned well with changes in the animal’s running speed and is strongly and linearly correlated to the logarithm of the animal’s running speed. Intriguingly, the cholinergic modulation remains strongly and linearly correlated to the speed of the animal’s neck movements during periods of stationary activity. Furthermore, we show that cholinergic modulation is unaltered during darkness. Lastly, we identify rearing, a discrete behavior where the mouse stands on its hindlimbs to scan the environment from an elevated perspective, is associated with higher cholinergic activity than expected from neck movements on the horizontal plane alone. Taken together, these data show that temporal dynamics in the cholinergic modulation of hippocampal circuits are fast enough to provide a potential running speed signal in real-time. Moreover, the data show that cholinergic modulation is primarily a function of the logarithm of the animal’s movement speed, both during locomotion and during stationary activity, with no significant interaction with visual inputs. These data advance our understanding of temporal dynamics in cholinergic modulation of hippocampal circuits and their functions in the context of neural coding of location and running speed.

## INTRODUCTION

Acetylcholine is an important neuromodulator of cognitive functions and behavior **(Hasselmo, 2006; Picciotto et al., 2012)** that are related to the processing of sensory information, memory, and navigation of physical and abstract mental spaces **(Solari and Hangya, 2018)**. Conversely, cholinergic dysfunctions are a hallmark of many neurological and psychiatric diseases that affect memory and the coding of spatial information and sensory information, such as Alzheimer’s disease **(Davies and Maloney, 1976; Hampel et al., 2018)**, autism spectrum disorders **(Karvat and Kimchi, 2014)**, schizophrenia **(Higley and Picciotto, 2014)**, and depression **(Higley and Picciotto, 2014)**. Experimental data from animal models demonstrate a role of acetylcholine in modulating learning and memory **(Hasselmo, 2006)**, attention to sensory stimuli **(Hasselmo and McGaughy, 2004; Yu and Dayan, 2005; Pinto et al., 2013)**, visual cue detection **(Gritton et al., 2016)**, and brain state transitions between waking, sleep, or behavioral states **(Marrosu et al., 1995; Xu et al., 2015; Harrison et al., 2016)**. In the hippocampal formation, cholinergic modulation of network dynamics, synaptic plasticity, and neuronal excitability supports the formation of spatial memories and cognitive functions supporting memory-guided navigation **(Blokland et al., 1992; Ohno et al., 1993, 1994; Stancampiano et al., 1999; Dannenberg et al., 2017)**. In particular, experimental data **(Rogers and Kesner, 2003)** and computational models **(Hasselmo, 1999, 2006)** propose an important role of acetylcholine in separating the encoding and retrieval of memory traces. The primary and major source of cholinergic innervation of the hippocampal formation is provided by cholinergic projection neurons in the medial septum/diagonal band of Broca (MSDB) **(Mesulam et al., 1983; Rye et al., 1984)**. MSDB cholinergic projection neurons have a key function in modulating hippocampal activity via a direct and an indirect pathway **(Alreja et al., 2000; Wu et al., 2000, 2003; Dannenberg et al., 2015)**. The indirect pathway is particularly important for modulating theta (6—10 Hz) rhythmic activity in the hippocampal local field potential via the modulation of glutamatergic and GABAergic projection neurons within the MSDB. Both the frequency and amplitude of theta rhythmic activity are correlated with the running speed of an animal **(Whishaw and Vanderwolf, 1973)**, and manipulations that disrupt either the power of theta **(Brandon et al., 2011; Koenig et al., 2011)** or the relationship to running speed of theta rhythmic frequency **(Winter et al., 2015; Dannenberg et al., 2020)** disrupt the coding of location by grid cells in the medial entorhinal cortex. Models of memory-guided navigation therefore propose a role of cholinergic modulation in the coding of location and running speed in the hippocampal formation **(Dannenberg et al., 2016)**. In fact, cholinergic activity has recently been demonstrated to be correlated with running speed in mice **(Jing et al., 2020; Zhang et al., 2021)**. However, the temporal dynamics in cholinergic activity and how they relate to changes in running speed and behavioral activities remain elusive, despite an ongoing debate in the field about the role of slow vs. fast time scales in cholinergic modulation **(Disney and Higley, 2020; Sarter and Lustig, 2020)**. To test the hypothesis that cholinergic activity changes fast enough to serve as a potential code for running speed in the hippocampal formation, we used a fiber-photometric approach **(Gunaydin et al., 2014; Lerner et al., 2015)** to monitor the population activity of cholinergic projection neurons in the MSDB of freely behaving mice. We next tested whether the activity of the septo-hippocampal cholinergic system is a function of characteristic behaviors, including stationary activities such as grooming and rearing. Lastly, we analyzed whether the observed temporal dynamics in cholinergic activity were a function of visual cues. The presented results demonstrate that the activity of the septo-hippocampal cholinergic system i) changes fast enough to match changes in running speed, ii) is strongly and linearly correlated to the logarithm of the animal’s running speed, iii) remains strongly correlated to the animal’s neck movements during stationary activity, iv) is elevated during rearing, and v) remains unchanged during darkness.

## MATERIALS AND METHODS

### Animals

Before surgery, mice were habituated to the experimenter and testing room. All experimental procedures were approved by the Institutional Animal Care and Use Committee for the Charles River Campus at Boston University. Mice were purchased from The Jackson Laboratory (Wildtype, C57Bl/6J; ChAT-IRES-Cre, B6;129S6-Chat^tm2(cre)Lowl^/J). Transgenic mice were maintained as homozygous, and heterozygous mice were used for experiments. For data collection, adult mice were housed in Plexiglas cages together with their siblings prior to surgery, but separated for individual housing after surgery, and maintained on a reversed 12-h light/12-h dark cycle. The housing cages contained a spherical treadmill that provided mice the opportunity to exercise.

### Viral Transduction and Light Fiber Implantation

Mice were injected with buprenorphine (0.1-mg/kg, s.c.) and atropine (0.1-mg/kg, i.p.), and survival surgery was performed under isoflurane anesthesia for virus injection and light fiber implantation targeting the MSDB. Four anchoring screws were positioned across the skull. For cell-type specific targeting of cholinergic MSDB neurons, we used stereotactically targeted virus injections of rAAV S1 FLEX-CAG-jGCaMP7s-WPRE (Lot v28549, Addgene, #104495 AAV-1) into the MSDB of 3— 6 months old ChAT-Cre mice. 2 × 250-nl virus solution was injected at two ventral sites within the MSDB. To that end, a craniotomy was performed 1.2-mm anterior and 0.7-mm lateral to Bregma, and the injection needle was lowered 4.8 and 4.4-mm at a 10° polar and −90° azimuth angle, following stereotactic coordinates from **Paxinos and Franklin (2008)**. The injection needle (34-g, beveled, WPI) was left in place for 3 and 5 min after the first and second injections (100-nl/min, UMP3 electrical pump, WPI) to prevent backflow of the injected virus solution. After virus injection was complete, the same opening in the skull was used to implant an implantable light fiber (200-µm core, N.A. 0.48, MFC_200/230-0.48_8mm_SMR_FLT, Doric lenses), 1.2-mm anterior and 0.7-mm lateral to Bregma. The light fiber was lowered 4.2-mm from the brain surface at an 10° polar and -90° azimuth angle and cemented on the animal’s skull with dental acrylic that was blackened by mixing in graphite. Animals were given buprenorphine (0.1-mg/kg, i.p.), enrofloxacin (7.5-mg/kg, i.p.), and ketoprofen (3-mg/kg, i.p.) during a 5-day postsurgical care period and allowed one week in total to fully recover after surgery before beginning of recordings.

### Behavioral Testing

All recordings were performed while animals foraged for small pieces of Froot Loops (Kellog Company, Battle Creek, MI, USA) in the open-field environment. A typical recording session lasted between 5 and 15-min. On a typical day, four recording sessions were performed. The first and last ones were performed during standard room lighting, the second and third recording sessions were recorded during darkness. A rectangular housing cage (34-cm x 28-cm) with transparent 20-cm high walls (n = 1 mouse) or an acrylic box (40-cm x 40-cm) with black 30-cm high walls and a visual cue card (n = 4 mice) were used as open field environments. The recording room contained no windows and was shielded with a laser-proof black curtain. Standard lighting conditions were performed with the room ceiling light turned on, and recordings during darkness were performed with the ceiling light turned off.

### Fiber Photometry System

We used a custom-built fiber photometry system inspired by **(Gunaydin et al., 2014; Lerner et al., 2015)**. The fiber photometry system used a 473-nm Omnicron Luxx laser modulated at 211-Hz and a 405-nm UV LED (Thorlabs, M405FP1) modulated at 531-Hz using optical choppers. Excitation at 405-nm served as an isosbestic control signal. Laser light was delivered into the brain via a system of fiber patch cords (Thorlabs) and a rotary joint (FRJ_1×1_FC-FC, Doric lenses). The laser light power entering the implanted light fiber was measured before and after every recording session and adjusted before the recording session to yield an estimate of 40—60 µW laser power delivered into the MSDB. Data were acquired at a sampling rate of 1-kHz.

### Video Tracking

Mice were video-tracked using a thermal camera (FLIR, SC8000) at a video frame rate of 30 frames/s controlled via TTL pulses generated by an external signal generator. Temperature values were color-coded with a gray scale and exported in mpeg-4 video format. TTL pulses sent from the camera were recorded along with the fiber-photometry system to synchronize the videos with the fiber-photometry signal.

### Analysis of Fiber Photometry Data

Data were re-sampled to match the 30-Hz video frame rate used for video-tracking of animal pose and behavior. The fiber-photometry signal was then processed to adjust for i) photobleaching that results in an exponential decay of the signal, and ii) motion artefacts. This was done in the following way. First, we subtracted the isosbestic control signal from the jGCaMP7s signal and fitted a 2^nd^-degree polynomial curve to that result. Next, we added the fitted curve to the isosbestic control signal, resulting in an adjusted control signal. We then formulated an optimization problem to find the two parameters α and β that minimized the term ∑(*s* * (*c* − *α* + *β*))^2^ where *s* stands for the jGCaMP7s signal, and *c* stands for the adjusted control signal. As a final step, we computed ΔF/F as 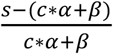. For analyses that average or compare data across sessions, the z-scored ΔF/F was used to account for variations in signal strength across sessions.

### Markerless Pose Estimation and Calculation of Running Speed

We used the deep learning tool DeepLabCut **(Mathis et al., 2018)** for markerless pose estimation of the animal’s neck, nose, left ear, right ear, tail base, and tail tip. The neck position was used to estimate the animal’s movement speed. For later identification of behavioral motifs using variational animal motion embedding (VAME), we labeled the animal’s neck, nose, left ear, right ear, three points along the spine, tail base, two points along the tail, and the tail tip.

### Variational Animal Motion Embedding (VAME)

To identify behavioral motifs and larger clusters of behavioral motifs (“communities”), we used the deep learning tool Variational Animal Motion Embedding (VAME) **(Luxem et al., 2022)**. All sessions that were included in this analysis were performed with the exact same camera setup and settings.

### Analysis Software and Code

Data analysis was performed using Matlab (MathWorks) and custom-written Matlab scripts. Code will be made available upon request.

## RESULTS

### The activity of cholinergic neurons is linearly correlated with the logarithm of the animal’s running speed

We used a recombinant adeno-associated virus (rAAV) and the Cre-loxP system to target the expression of the genetically encoded fluorescent Calcium indicator jGCaMP7s **(Dana et al., 2019)** to the cholinergic subpopulation in the MSDB of transgenic mice that express the Cre-recombinase under the control of the choline-acetyltransferase (ChAT) promoter (ChAT-Cre mice). This allowed us to monitor the population activity of cholinergic neurons in the MSDB of a total of five adult ChAT-Cre mice (two females, three males; **Table 1**) at high temporal resolution during free foraging in an open field environment. The temporal resolution of measurements in cholinergic activity was only constrained by the kinetics of the Calcium sensor jGCaMP7s that has a reported half-decay time of 1.69-s **(Dana et al., 2019)**, providing sufficient temporal resolution to analyze the temporal dynamics in cholinergic activity as a function of changes in the running speed of freely foraging mice. To measure the running speed of mice, we used a thermal camera (FLIR SC8000) positioned above the center of the open field box. We used the markerless pose estimation tool DeepLabCut **(Mathis et al., 2018)** to identify, in each video frame, the neck position of the mouse which was used to compute the running speed **(Figure 1G)**. Consistent with a recent study using fiber photometry to monitor the release of acetylcholine in the hippocampus in freely moving mice **(Zhang et al., 2021)**, we found that the population activity of cholinergic neurons in the MSDB was correlated to the running speed during periods of locomotion **(Figure 1A)**. However, we were intrigued by the fact that the cholinergic activity showed the largest fluctuations at low running speeds **(Figure 1B)** resulting in speed tuning curves of cholinergic activity that are best fitted by a saturating exponential function, both in single sessions (see **Figure 1B** for one example) and when averaging data across 103 sessions from five mice **(Figure 1C)**. Since the temporal dynamics in the cholinergic activity were not reflected in the speed tuning curves and showed larger variations at lower running speeds, we asked whether changes in cholinergic activity may correlate with changes in the *logarithm* of the running speed. In fact, cholinergic activity showed a strong and strikingly linear correlation to the logarithm of the running speed **(Figures 1D—F)**. Theoretical work has been shown that information from optical flow could contribute to coding of location and running speed in the medial entorhinal cortex **(Raudies et al., 2012, 2016)**. Experimental data showing the contribution of visual inputs on the spatial accuracy of grid cell firing in the medial entorhinal cortex **(Chen et al., 2016; Pérez-Escobar et al., 2016; Dannenberg et al., 2020)** support the idea that movement information from optical flow can contribute to speed coding in the hippocampal formation. We therefore tested the hypothesis that the relationship between the activity of cholinergic neurons in the MSDB and the animal’s running speed is changed in the absence of visual information by testing the effect of darkness on the speed tuning of cholinergic activity using a linear mixed effects model of data obtained from recordings under standard lighting conditions and darkness. Interestingly, we found no substantial change in the speed tuning of cholinergic activity during darkness **(Figures 1H and 1I; Table 2)**. These data indicate that the temporal dynamics in the activity of cholinergic neurons in the MSDB are primarily a function of the animal’s movement speed with a negligible influence of visual cues.

**Table 1.** Number of sessions and time span of recordings per mouse. Total number of sessions was 103 from 5 mice recorded over a time span of 10—15 days per mouse.

| Mouse ID | Sex | Number of sessions | Time span of recordings (days) |
| --- | --- | --- | --- |
| 611915 | Female | 20 | 12 |
| 611916 | Female | 46 | 16 |
| 611926 | Male | 8 | 10 |
| 616195 | Male | 17 | 13 |
| 616211 | Male | 12 | 15 |

**Figure 1.**
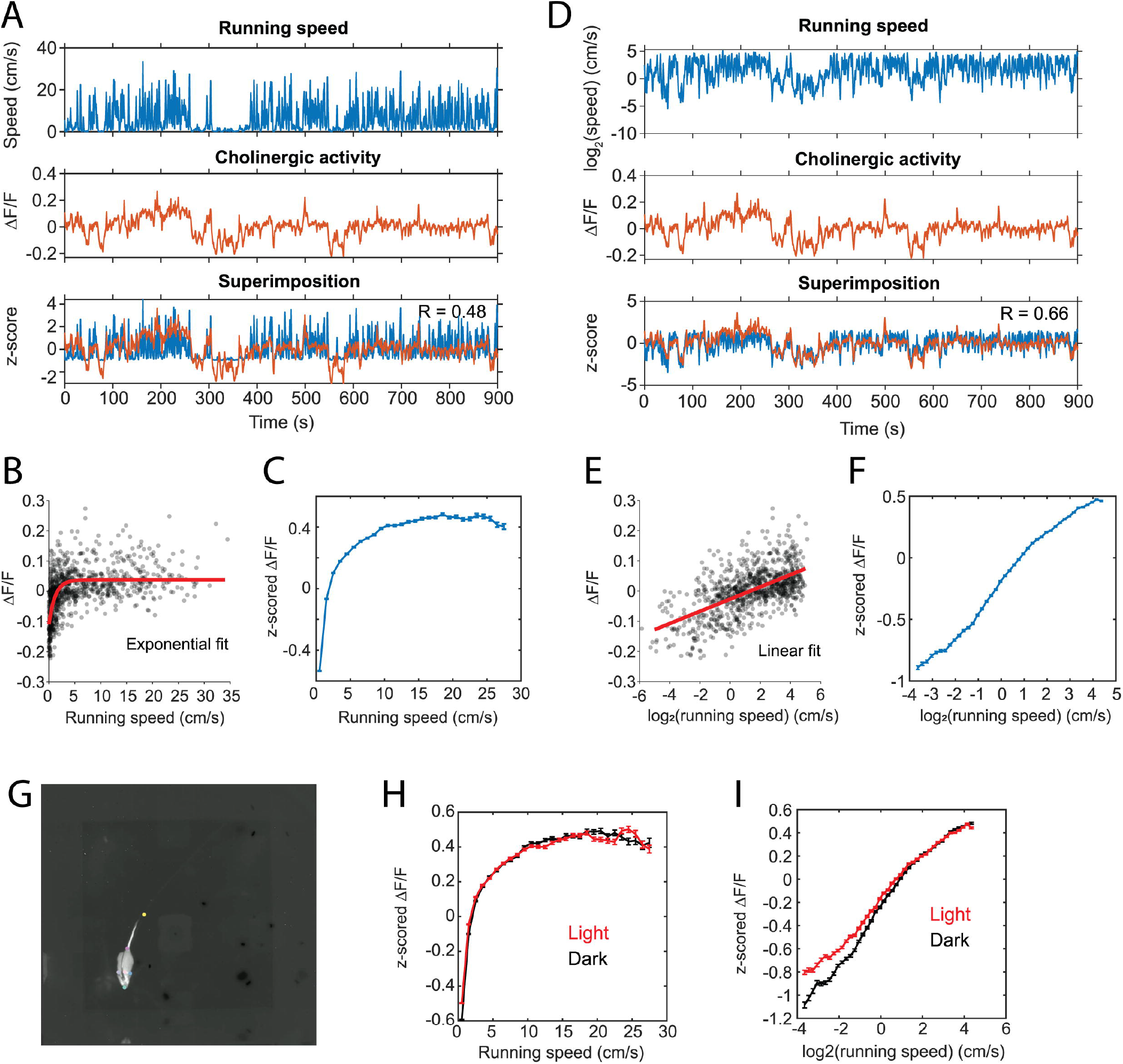
Linear correlation between the population activity of MSDB cholinergic neurons and the logarithm of an animal’s running speed. **(A)** Data on running speed and activity of cholinergic neurons in the MSDB measured via fiber-photometry for one example session. Note the high fluctuations in the fiber photometry signal during low running speeds. **(B)** Scatter plot of cholinergic activity vs. running speed; data points were sampled in 1-s intervals from the time series data shown in (A). Red line shows the exponential fit to the data. **(C)** Speed tuning curve of cholinergic activity. Data show mean ± s.e.m. of speed-binned data with a bin width of 1-cm/s; data from 103 sessions recorded from 5 mice. **(D)** Same data as in (A) but now plotting the logarithm of running speed. **(E)** Scatter plot of cholinergic activity vs. the logarithm of the animal’s running speed; data points were sampled in 1-s intervals from the time series data shown in (D). Red line shows the linear fit to the data. **(F)** Tuning curve shows cholinergic activity as a function of the logarithm of running speed; data show mean ± s.e.m. of binned data with a bin width of 0.1; data from 103 sessions recorded from 5 mice; **(G)** One video frame showing the DeepLabCut labels for neck, nose, left ear, right ear, tail base, and tail tip. Gray scale of the video shows the temperature of the mouse recorded via a thermal camera. **(H)** Speed tuning curves of cholinergic activity comparing data from light (red, n = 55) and darkness (black, n = 47) sessions; data presented as in (C) **(I)** Data presented as in (F) comparing data from light (red, n = 55) and darkness (black, n = 47) sessions; no significant difference between light and darkness sessions was found; see Table 2 for statistics. R = Pearson’s correlation coefficient.

**Table 2.** Results of a linear mixed-effects model of the z-scored ΔF/F of cholinergic activity with three fixed effects, namely the y-intercept, the logarithm of allocentric neck movement speed, and the interaction between the logarithm of allocentric neck movement speed and darkness. Random effects of the animal (n = 5) and sessions (n = 102, 55 “light” sessions and 47 “dark” sessions) on all fixed effects were included in addition to a random error term. Total number of observations: 1310717; Fixed effects coefficients: 3; Random effects coefficients: 321; Covariance parameters: 7. SE = standard error; DF = degrees of freedom; CI = confidence interval.

| Fixed effects coefficients |  |  |  |  |  |  |
| --- | --- | --- | --- | --- | --- | --- |
| Name | Estimate | SE | t-statistic | DF | p-value | 95% CI |
| y-intercept | -0.169 | 0.072 | -2.35 | $1.31 \times 10^6$ | 0.0186 | [-0.310 -0.028] |
| $\log_2(\text{speed})$ | 0.221 | 0.019 | 11.92 | $1.31 \times 10^6$ | <0.0001 | [0.185 0.258] |
| $\log_2(\text{speed})^*$<br>darkness | 0.013 | 0.013 | 0.99 | $1.31 \times 10^6$ | 0.3213 | [-0.013 -0.038] |
| Model statistics |  |  |  |  |  |  |
| AIC: $3.424 \times 10^6$ BIC: $3.424 \times 10^6$ Log-likelihood: $-1.7 \times 10^6$ Deviance: $3.424 \times 10^6$ | | | | | | |

### The activity of cholinergic neurons is linearly correlated to neck movements during stationary behavior

We noticed that cholinergic activity showed huge variations for running speeds below 5—10 cm/s. Our previous analysis primarily focused on the analysis of locomotion as opposed to other characteristic behaviors or body movements that are not related to locomotion but can also occur during stationary periods, i.e. in the absence of locomotion. However, the observed large variance in cholinergic activity for small running speeds below 3-cm/s and the fact that the observed linear correlation of cholinergic activity with the logarithm of the animal’s running speed holds true even for very small running speeds suggest that cholinergic activity might also be correlated to neck movements while the mouse is stationary in space **(Figure 1F)**. To make sure that our analysis of neck movements during stationary periods was not confounded by the effects of stop-and-go-patterns during locomotion, we analyzed a subset of n = 16 out of 103 sessions in which the mouse was stationary (allocentric neck movement speed < 3-cm/s) for more than two thirds of the total length of the recording session **(Figure 2)**. Intriguingly, we found that the strong and linear correlation of cholinergic activity with the logarithm of the animal’s running speed extended to the logarithm of the animals’ neck movement speed during stationary activity. To test whether the speed tuning of cholinergic activity during stationary activity depends on visual cues or optic flow, we again modeled the effect of darkness on speed tuning during stationary activity using a linear mixed-effects model **(Table 3)**. As for the speed tuning during locomotion, we found no significant effect of darkness on speed tuning of cholinergic modulation during stationary activity. In summary, these data suggest that a potential speed code by the population activity of cholinergic MSDB neurons does not distinguish between stationary periods and locomotion but instead codes for the whole range of translational neck movement speeds in the horizontal plane between 0-cm/s and the animal’s maximal running speed.

**Figure 2.**
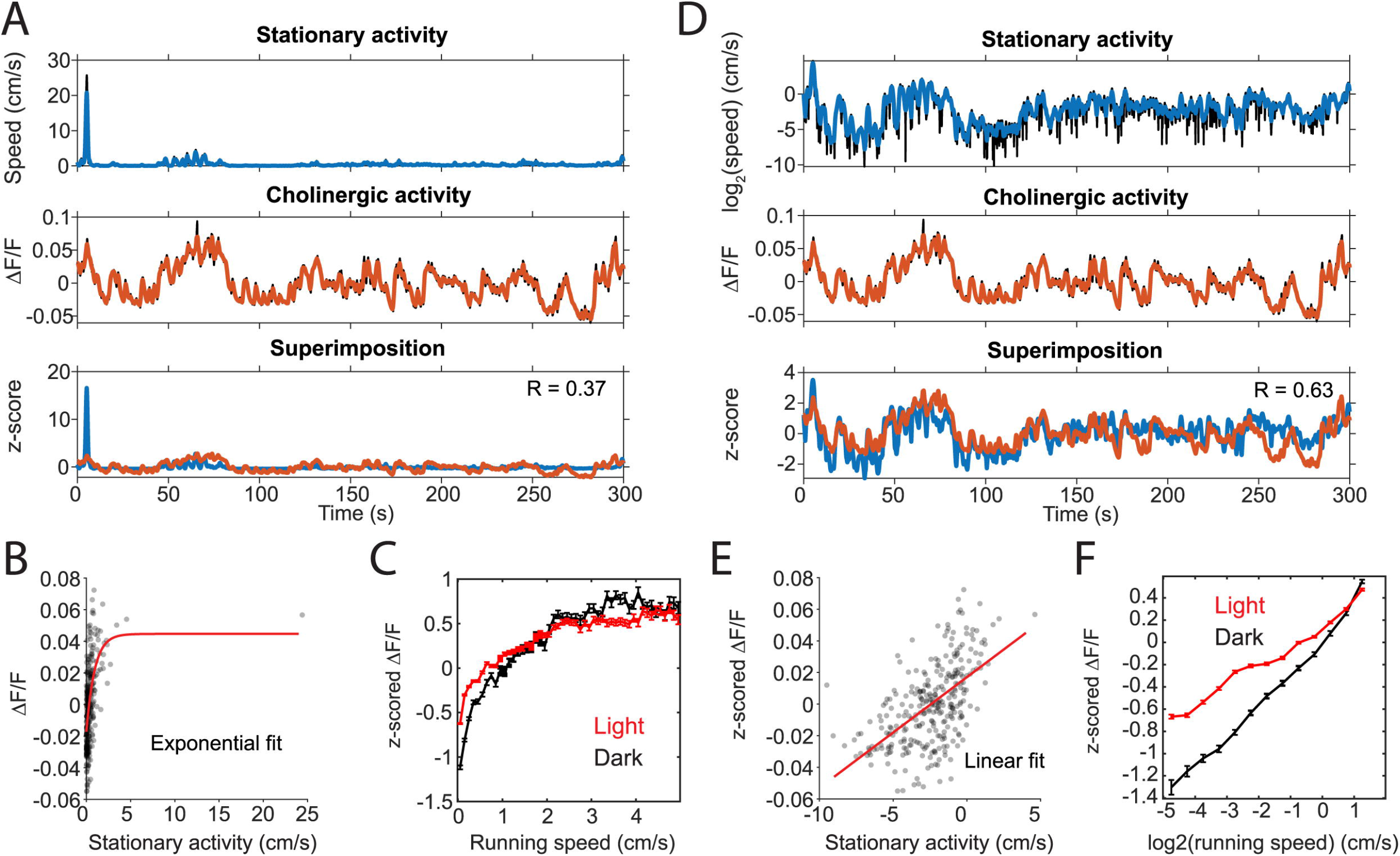
The activity of cholinergic neurons is linearly correlated to neck movements during stationary behavior. **(A)** Data on the speed of neck movements during stationary behaviors and activity of cholinergic neurons in the MSDB measured via fiber-photometry for one example session, in which the mouse remained stationary (running speed < 3-cm/s) for more than two thirds of the session length. Note the high fluctuations in the fiber photometry signal during stationary activity. **(B)** Scatter plot of cholinergic activity vs. running speed; data points were sampled in 1-s intervals from the time series data shown in (A). Red line shows the exponential fit to the data. **(C)** Speed tuning curves of cholinergic activity comparing data from light (red, data from 13 sessions, 3 mice) and darkness (black, data from 3 sessions, 2 mice) sessions. Data show mean ± s.e.m. of speed-binned data with a bin width of 1-cm/s. **(D)** Same data as in (A) but now plotting the logarithm of neck movement speed. **(E)** Scatter plot of cholinergic activity vs. the logarithm of the animal’s neck movement speed; data points were sampled in 1-s intervals from the time series data shown in (D). Red line shows the linear fit to the data. **(F)** Tuning curve shows cholinergic activity as a function of the logarithm of neck movement speed; data show mean ± s.e.m. of binned data with a bin width of 0.1; data on light sessions (red) from 13 sessions, 3 mice; data on darkness sessions (black) from 3 sessions, 2 mice; no significant difference between light and darkness sessions was found; see Table 3 for statistics. R = Pearson’s correlation coefficient.

**Table 3.** Results of a linear mixed-effects model of the z-scored ΔF/F of cholinergic activity with three fixed effects, namely the y-intercept, the logarithm of allocentric neck movement speed, and the interaction between the logarithm of allocentric neck movement speed and darkness. Random effects of the animal (n = 3) and sessions (n = 16, 13 “light” sessions and 3 “dark” sessions) on all fixed effects were included in addition to a random error term. Total number of observations: 146234; Fixed effects coefficients: 3; Random effects coefficients: 57; Covariance parameters: 7. SE = standard error; DF = degree of freedom; CI = confidence interval.

| **Fixed effects coefficients** |  |  |  |  |  |  |
| --- | --- | --- | --- | --- | --- | --- |
| Name | Estimate | SE | t-statistic | DF | p-value | 95% CI |
| y-intercept | 0.232 | 0.127 | 1.83 | $1.46 \times 10^5$ | 0.067 | [-0.016 0.481] |
| $\log_2(\text{speed})$ | 0.226 | 0.028 | 8.14 | $1.46 \times 10^5$ | <0.0001 | [0.171 0.280] |
| $\log_2(\text{speed})^*$<br>darkness | 0.068 | 0.050 | 1.37 | $1.46 \times 10^5$ | 0.170 | [-0.029 0.166] |
| **Model statistics** |  |  |  |  |  |  |
| AIC: $3.597 \times 10^5$ BIC: $3.598 \times 10^5$ Log-likelihood: $-1.8 \times 10^5$ Deviance: $3.597 \times 10^5$ | | | | | | |

### Temporal dynamics in cholinergic activity are fast enough to match changes in movement speed

Speed tuning curves of cholinergic activity do not provide any information about the temporal dynamics in cholinergic modulation. If changes in the population activity of cholinergic neurons are fast enough to align with changes in the speed of neck movements during stationary activity or running, the population activity of cholinergic neurons could potentially be used in computational models to provide a movement speed signal in real-time. To quantify the temporal dynamics in cholinergic modulation, we computed the integration time window for cholinergic activity that maximized the correlation between cholinergic activity and the logarithm of the neck movement speed of an animal. When averaging across 103 sessions from five mice, we found that an integration time window of just 1.3-s optimized the correlation between the population activity of cholinergic MSDB neurons and the animals’ running speed during locomotion or neck movement speeds during stationary activity **(Figure 3A)**. Notably, 1.3-s is shorter than the reported half-decay time of jGCaMP7s, the fluorescent Calcium sensor used in this study to monitor the activity of cholinergic MSDB neurons, indicating that our time scale estimate of changes in cholinergic activity reached the temporal resolution limit of what we could detect in this study. We conclude that changes in MSDB cholinergic activity are fast enough to match changes in running speed during locomotion or neck movements during stationary behavior and that changes in cholinergic activity could potentially be even faster than we were able to detect in this study. Optimal integration time windows that maximize the correlation of the smoothed cholinergic activity and the neck movement speed of the animal were computed for each session. This allowed us to analyze the distribution of optimal integration windows across sessions **(Figure 3B)**. We found that the distribution is quite narrow with a strong peak between 0.8-s and 1-s, further validating our previous finding that changes in cholinergic activity are fast and align well with the time scale of changes in the animal’s running speed during locomotion or neck movement speeds during stationary behavior. We next tested whether the temporal alignment of cholinergic activity with locomotion or neck movements during stationary activity depends on visual inputs or optic flow by comparing data during standard room lighting and darkness sessions (light: n = 55 sessions, five mice; dark: n = 47 sessions, four mice). Consistent with the results of our previous analysis of speed tuning curves of cholinergic activity during darkness, we found that the relationship between changes in cholinergic activity and changes in the animal’s running speed or neck movement speeds during stationary activity remained unaltered during darkness **(Figures 3C and 3D)**; there was no significant difference between the Pearson’s correlation coefficients for the correlation between 1.3-s-smoothed cholinergic activity and the logarithm of movement speed between sessions recorded during standard lighting and darkness (R_Light_ = 0.46 ± 0.17; R_Dark_ = 0.46 ± 0.13; t(100) = -0.192, p = 0.85, 95% CI = [-0.07 0.05]). Likewise, the distribution of optimal integration windows computed for each session were not significantly different between standard lighting and darkness conditions (p = 0.29, Kolmogorov-Smirnov test). Lastly, we asked whether the temporal dynamics in cholinergic activity during stationary activity align with changes in the speed of an animal’s neck movements on a similarly fast time scale as they do with changes in running speed during locomotion. To answer this question, we analyzed the subset of n = 16 sessions in which the mice remained stationary (allocentric neck movement speed < 3-cm/s) for more than two thirds of the session time. We found that the integration time window that maximizes the correlation of cholinergic activity and the neck movement speed during stationary activities was 1.7-s, thus not substantially different from the integration time window of 1.3-s that maximized the correlation of cholinergic activity in the complete data set **(Figures 3E and 3F)**.

**Figure 3.**
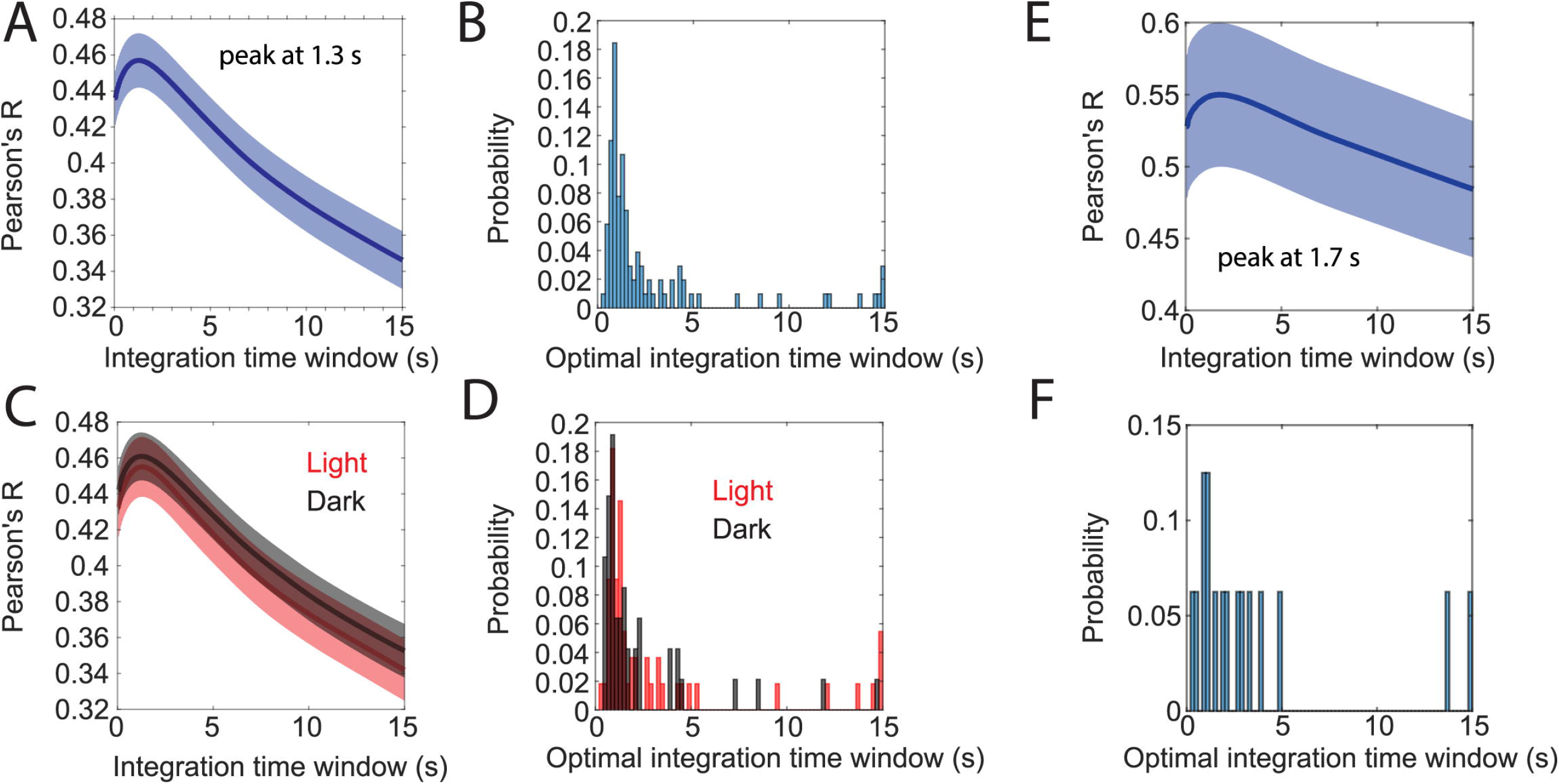
Temporal dynamics in cholinergic activity align with changes in movement speed. **(A)** Data show Pearson’s correlation coefficients between moving averages of cholinergic activity and the animal’s movement speed as a function of the time window used for computing the moving average of cholinergic activity. Blue line and shaded area show the mean ± s.e.m. of data from n = 103 sessions from five mice. The average peaks at 1.3-s. **(B)** Distribution of the optimal time windows, computed for each session, that maximize the correlation between moving averages of cholinergic activity and movement speed. The histogram shows a peak between 0.8-s and 1-s. **(C, D)** Data presented as in (A) and (B) but comparing data from light (red, n = 55 sessions, five mice) and darkness sessions (black, n = 47, 4 mice). No significant difference was found between light and darkness sessions shown in (C) when comparing values at 1.3-s, t(100) = -0.192, p = 0.85. **(E)** Subset of data presented in (A) that only includes n = 16 sessions from three mice, in which the mice spent at least two thirds of the session time engaged in stationary behaviors (running speed < 3-cm/s). The average peaks at 1.7-s. **(F)** Same subset of data on stationary activity as shown in (E). Histogram shows the distribution of the optimal time windows, computed for each session, that maximize the correlation between moving averages of cholinergic activity and neck movement speed during stationary activity.

### Cholinergic neurons in the MSDB are more active during rearing

The data presented so far indicate a strong and linear correlation of cholinergic MSDB activity with the neck movement of the animal, regardless whether those movements are caused by translational movements due to walking or running or by characteristic behaviors such as grooming or rearing. However, a recent study has shown that cholinergic activity is low during the occurrence of sharp-wave-ripples in the hippocampus, 40—100 ms long events of synchronous network activity detected in the local field potential that typically occur while the mouse is stationary **(Zhang et al., 2021)**. Conversely, rearing, a discrete behavior that is associated with attention, exploration, and integration of sensory inputs, has been shown to be accompanied by an increase in the frequency of theta rhythmic activity and theta-gamma coupling **(Barth et al., 2018)**, hippocampal network states that are typically associated with higher cholinergic activity. We therefore asked whether characteristic behaviors affect the activity of cholinergic MSDB neurons. To answer this question, we first chose an unbiased approach to identify discrete behavioral motifs using the deep learning tool “VAME” (variational animal motion embedding) **(Luxem et al., 2022)**. VAME takes the body part positions identified by DeepLabCut as input to train a model of animal motion that is then used to classify distinct motion patterns into clusters of similar discrete behavioral motifs. To improve the accuracy of VAME, we increased the number of DeepLabCut labels from six to eleven and relabeled the videos from n = 74 sessions of three mice where the videos were taken with the same camera and video settings to ensure robust model performance across sessions. VAME identified four main clusters (“communities”) of behavioral motifs **(Figures 4A and 4B)** that could be identified post-hoc by the experimenter as “exploratory running”, “exploratory walking”, “grooming”, and “rearing”. On average, mice spent 1.4% of their time grooming, 26.2% of their time walking, 70.4% of their time running, and 2.1% of their time rearing.

**Figure 4.**
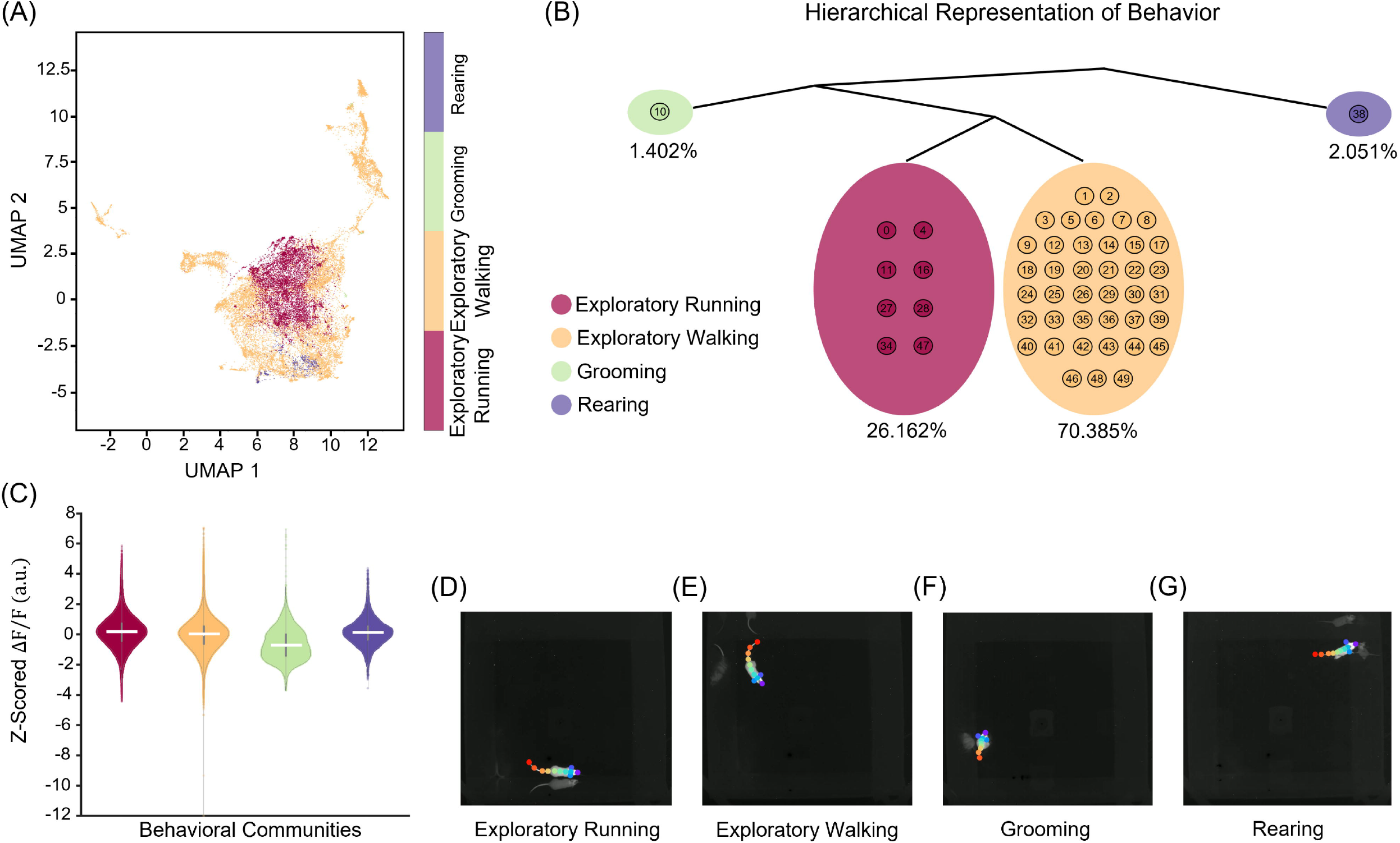
Variational Animal Motion Embedding (VAME) reveals four distinct behaviors during open field exploration. **(A)** Unifold Manifold Approximation (UMAP) embedding of the latent representation encoded from VAME’s recurrent neural network, color-coded based on four human expert-labeled behavioral communities corresponding to exploratory running, exploratory walking, grooming, and rearing. **(B)** Hierarchical representation of the four communities of behavioral motifs in (A). Percentages below each community highlight the time duration observed for that behavior across a total of ten hours of open field exploration (n = 74 sessions). **(C)** Z-scored ΔF/F associated with each behavioral community. Horizontal solid white lines indicate the medians. **(D-G)** Representative video frames for each behavioral community identified with VAME.

To test whether those four identified main behavioral clusters influenced the activity of cholinergic neurons in the MSDB, we investigated the effects of behavioral clusters on cholinergic activity using a linear mixed effects model. As expected, we found a highly significant effect of the logarithm of the neck movement speed but no significant effect on darkness on cholinergic activity. However, we found a significant positive effect of rearing on cholinergic activity, while exploratory walking, exploratory running, and grooming had no significant impact on the activity of cholinergic neurons **(Figure 4C and Table 4; Supplementary Figure S1)**.

**Table 4.**
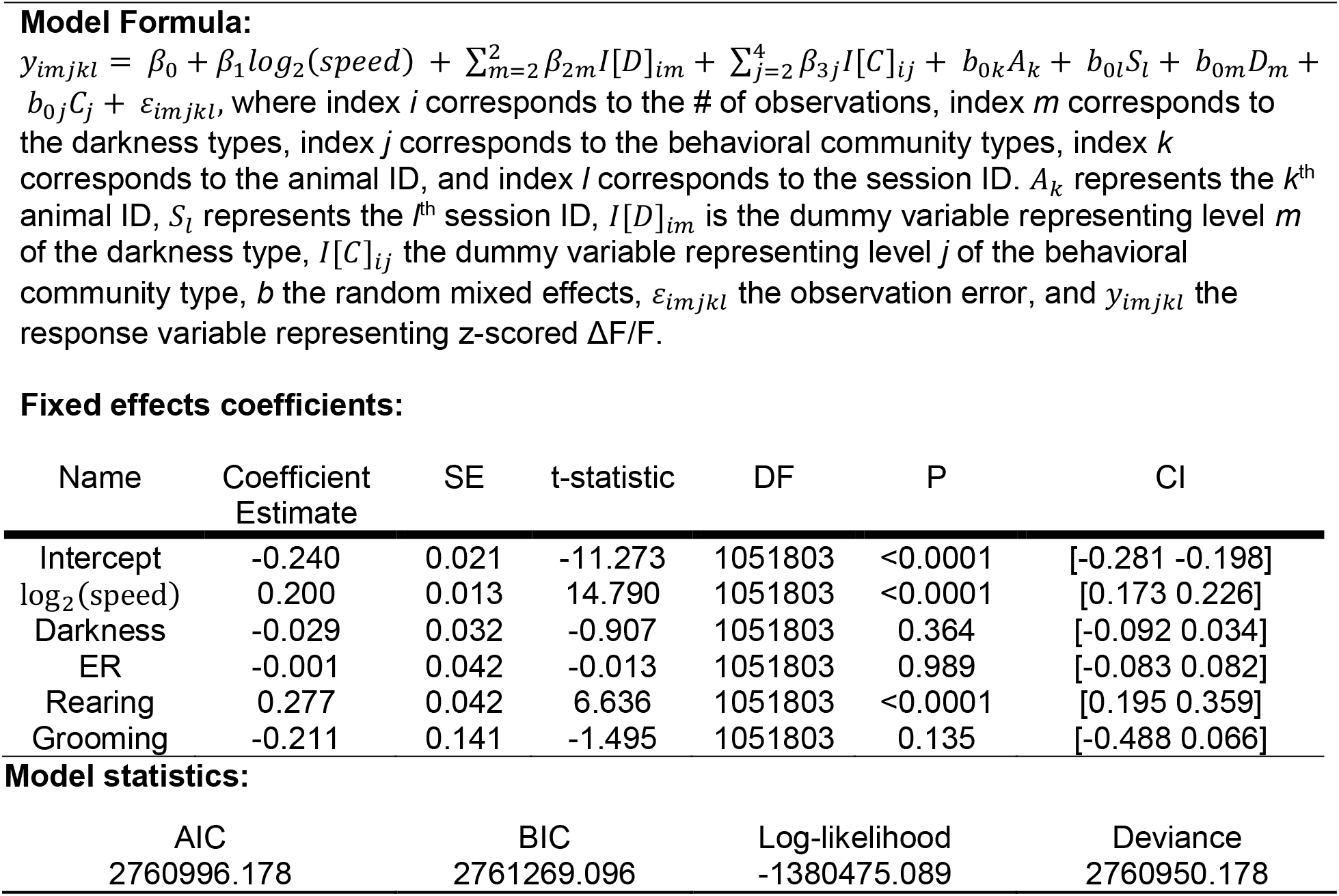
Results of a linear mixed-effects model of the z-scored ΔF/F of cholinergic activity with six fixed effects, which include y-intercept, the logarithm of allocentric neck movement speed, darkness, and the behavioral communities as identified through VAME. Random effects of the animal (n = 3) and sessions (n = 41 light and n = 32 dark) on all fixed effects were included in addition to a random error term. Total number of observations: 1051803; fixed effects coefficients: 6; random effects coefficients: 380; covariance parameters: 17. ER, Exploratory Running.

**Fixed effects coefficients:**
| Name | Coefficient Estimate | SE | t-statistic | DF | P | CI |
| --- | --- | --- | --- | --- | --- | --- |
| Intercept | -0.240 | 0.021 | -11.273 | 1051803 | <0.0001 | [-0.281 -0.198] |
| $\log_2(speed)$ | 0.200 | 0.013 | 14.790 | 1051803 | <0.0001 | [0.173 0.226] |
| Darkness | -0.029 | 0.032 | -0.907 | 1051803 | 0.364 | [-0.092 0.034] |
| ER | -0.001 | 0.042 | -0.013 | 1051803 | 0.989 | [-0.083 0.082] |
| Rearing | 0.277 | 0.042 | 6.636 | 1051803 | <0.0001 | [0.195 0.359] |
| Grooming | -0.211 | 0.141 | -1.495 | 1051803 | 0.135 | [-0.488 0.066] |

| AIC | BIC | Log-likelihood | Deviance |
| --- | --- | --- | --- |
| 2760996.178 | 2761269.096 | -1380475.089 | 2760950.178 |

Taken together, these data demonstrate that cholinergic neurons in the MSDB are more active during rearing as compared to other characteristic behaviors such as grooming, exploratory walking, or exploratory running.

## DISCUSSION

Experiments and analyses presented in this study investigate the temporal dynamics of cholinergic activity in the septo-hippocampal system in freely foraging mice as a function of the animal’s running speed during locomotion, the speed of neck movements during stationary activity, visual inputs, and discrete behavioral motifs identified by unsupervised variational animal motion embedding. The presented results demonstrate that the temporal dynamics in the population activity of MSDB cholinergic neurons are fast enough to align with the temporal dynamics of running speed during locomotion periods as well as temporal dynamics in the speed of neck movements during periods of stationary activity such as grooming or stationary head movements. Intriguingly, the logarithm of the speed of neck movements correlated strongly and linearly across the whole range of speed values between 0-cm/s and the maximal running speed with no detectable change in the speed tuning of cholinergic activity at the transition point from stationary activity to locomotion. Furthermore, the quantification of temporal dynamics in the cholinergic activity and their relationship to the speed of the animal revealed that cholinergic neurons can change their activity fast enough to match changes in running speed. Notably, no differences in cholinergic activity and its relationship to running speed or neck movements during stationary activity were detected during darkness. Lastly, an analysis of the effect of four discrete clusters of behavioral motifs that correspond to exploratory running, exploratory walking, grooming, and rearing revealed that rearing is associated with high cholinergic activity.

Importantly, cholinergic modulation can act on different time scales ranging from milliseconds to minutes (transient or tonic, fast or slow) with important consequences on cortical dynamics. However, the temporal dynamics in cholinergic activity, particularly in relation to running speed, have not been quantified so far, mostly due to a lack of recording techniques that allowed the recording of cholinergic activity in the MSDB at fast temporal resolution in freely behaving mice. Previous measurements of cholinergic activity in the septo-hippocampal system used microdialysis **(Marrosu et al., 1995)** or amperometry **(Parikh et al., 2007; Teles-Grilo Ruivo et al., 2017)** techniques to measure changes in the release of acetylcholine at a temporal resolution of minutes or multiple seconds, respectively. In this study, we used a fiber photometry approach to monitor the population activity of cholinergic neurons in the MSDB on a time scale of ∼1-s that proved fast enough to study the temporal dynamics in relation to changes in movement speed. Cholinergic projection neurons in the MSDB are the primary and major source of cholinergic innervation of the hippocampal formation. **(Mesulam et al., 1983; Rye et al., 1984)** and have a key function in modulating hippocampal activity via a direct or indirect pathway **(Alreja et al., 2000; Dannenberg et al., 2015)**. In addition to cholinergic neurons, the MSDB contains glutamatergic and GABAergic neurons. All three cell types form an interconnected network within the MSDB and project to the hippocampal formation **(Robinson et al., 2016; Müller and Remy, 2017)**. While the MSDB as a whole is important for modulating hippocampal network functions related to memory, spatial cognition, and memory-guided navigation, the population of cholinergic neurons in the MSDB is of particular interest because the neurotransmitter acetylcholine is an important neuromodulator of cognitive functions and behavior. Importantly, acetylcholine can modulate hippocampal network functions via binding to nicotinic and muscarinic receptors with different time courses of effect. Moreover, the activity of cholinergic MSDB neurons can modulate hippocampal network functions indirectly via recruiting glutamatergic and GABAergic MSDB projection neurons **(Alreja et al., 2000)**. Experimental data show that optogenetic activation of glutamatergic MSDB neurons initiated movements in mice **(Fuhrmann et al., 2015; Robinson et al., 2016)** and optogenetic activation of GABAergic neurons modulated theta rhythmic activity **(Dannenberg et al., 2015; Zutshi et al., 2018)** that is correlated to the running speed of an animal. Because of these cholinergic effects in the medial septum, fiber-photometric recordings of the activity of cholinergic projection neurons in the MSDB allows a more holistic interpretation of the temporal dynamics in cholinergic activity compared to data on the synaptic release of acetylcholine in the hippocampal formation. Future studies analyzing and comparing the temporal dynamics in synaptic release of acetylcholine in the hippocampal formation will need to determine whether there are differences in the temporal dynamics of the population activity of MSDB cholinergic neurons and the temporal dynamics in the synaptic release of acetylcholine in the hippocampus and related structures.

The fast temporal dynamics in conjunction with the strong and linear correlation of cholinergic activity with the logarithm of the animal’s running speed during locomotion support the hypothesis that the activity of cholinergic neurons in the MSDB can provide a speed signal to the hippocampus and medial entorhinal cortex. This interpretation is further supported by our findings that the temporal dynamics in cholinergic activity and the correlation to running speed are unaltered during darkness, suggesting that the provided speed signal could be used for path integration. Computational models of path integration typically rely on a linear speed signal. Speed cells in the medial entorhinal cortex have been suggested to provide a code for running speed **(Kropff et al., 2015; Hinman et al., 2016; Carvalho et al., 2020)**. However, the temporal dynamics in speed cells’ firing rates below 1 second have been shown to not accurately match changes in the running speed of animals **(Góis and Tort, 2018; Dannenberg et al., 2019)**. Data presented in this study suggest that cholinergic modulation could provide a speed signal on the time scale of 1 second used by a neural reader mechanism **(Buzsáki, 2010)** in the hippocampus and entorhinal cortex. Importantly, the signal is linearly correlated to the logarithm of the animal’s running speed and extends into stationary periods. The latter is a potential advantage over other proposed speed codes such as the speed code by entorhinal speed cells **(Kropff et al., 2015; Ye et al., 2018)** and a proposed speed signal represented by theta rhythmic activity in the local field potential or by theta rhythmic spiking of neurons **(Hinman et al., 2016)**. In previous studies, the speed signal represented by firing rates of neurons in the medial entorhinal cortex was only tested on running speeds above a speed threshold of 3-cm/s. Likewise, a potential speed code by theta frequency or theta amplitude is restricted to periods of locomotion due to the absence of theta rhythmic activity during most stationary activities such as grooming. However, the speed of head movements or the speed of changes in body position is likely an important factor for path integration, even if those movements occur while the mouse remains stationary.

These data provided experimental evidence that cholinergic MSDB neurons are an important component of neural circuits that control or code for running speed. Moreover, lesions of cholinergic projection neurons impair path integration **(Martin and Wallace, 2007; Martin et al., 2007; Hamlin et al., 2013; Yoder et al., 2017)**, providing further experimental support that cholinergic activity could be used as a speed signal by the hippocampus and medial entorhinal cortex. An alternative, not mutually exclusive, interpretation of the current data is that the activity of MSDB cholinergic neurons is primarily correlated to physical activity instead of to the logarithm of movement speed. This interpretation is supported by the fact that cholinergic activity remains strongly and linearly correlated to the logarithm of the speed of neck movements during stationary activity. Future experiments need to address questions related to the colinearity of movement speed and physical activity.

Acetylcholine release in the hippocampal formation is correlated to active exploration and can dramatically increase responses to stimuli in the visual cortex **(Niell and Stryker, 2010)**. Cholinergic activity is also significantly correlated with cue detection **(Gritton et al., 2016)** and visual perception **(Goard and Dan, 2009; Niell and Stryker, 2010; Pinto et al., 2013)** and attention **(Hasselmo and McGaughy, 2004)**, and cholinergic signaling is a hallmark in disorders of attention and cognitive control **(Wallace et al., 2011; Higley and Picciotto, 2014)**. These data suggested that temporal dynamics in cholinergic activity might be a function of visual inputs or optic flow. We therefore tested whether the observed temporal dynamics in cholinergic activity and their relationship to running speed changed during darkness. Interestingly, the relationship of cholinergic dynamics to changes in running speed remained unaltered during locomotion in darkness. Likewise, the relationship of cholinergic dynamics to changes in the speed of the animal’s neck movements during darkness appeared unaltered during stationary activity. These findings are consistent with findings of a previous study showing that glutamatergic cells in the MSDB remain tuned to running speed in the absence of optic flow **(Justus et al., 2016)**. The finding that visual inputs do not alter the temporal dynamics of cholinergic activity as a function of running speed are similar to findings in previous studies of a potential speed code by firing rate in neurons of the entorhinal cortex **(Dannenberg et al., 2020)** and posterior parietal cortex **(Alexander et al., 2022)** that demonstrated no change in the speed tuning of normalized firing rates. However, those studies showed a change in the *gain* of speed tuning of absolute firing rates. Since the analysis of fiber-photometry data requires the normalization of cholinergic activity, we cannot compare absolute values of cholinergic activity across sessions. It thus remains to be tested whether the absence of visual inputs can change the gain in the speed tuning of cholinergic activity.

Empirical data suggest that rearing, a behavioral action in a stationary location where mice stand on their hindlimbs to scan the environment from an elevated perspective, is associated with a distinct brain state supporting the encoding of spatial information from distant visual cues **(Barth et al., 2018)**. In fact, we found that rearing, was correlated with higher cholinergic activity. These data are consistent with the role of acetylcholine for processing of visual information and the encoding of memories **(Hasselmo, 2006)**. However, there is an alternative explanation that deserves to be considered. During rearing, the mouse moves vertically instead of horizontally. Due to the centered position of the camera used for the tracking of mice, vertical movements cannot be detected. Future studies need to address the question whether the observed increase in cholinergic activity during rearing is primarily due to the cognitive demand of processing visual information or encoding novel memories or due to coding the movement speed in the vertical axis.

## AUTHOR CONTRIBUTIONS

H.D. and M.E.H. designed the research. H.D., H.L., and P.N. conducted the experiments. H.D., J.D.K., and K.H. analyzed the data. H.D., J.D.K., and M.E.H. wrote and revised the manuscript.

## ACKNOWLEDGMENTS

We thank Dr. Kurtulus Golcuk and Dr. Heinz Beck for sharing their technical knowledge on constructing a custom-built fiber-photometry system.

## FIGURE LEGENDS

**Supplementary Figure S1. Differences in movement speed across behavioral clusters**. Violin plots show the distribution of movement speeds within behavioral clusters identified by VAME. Horizontal solid white lines indicate the medians.

