## Supplementary figures and images for "Temporal dynamics of cholinergic activity in the septo-hippocampal system"

### Supplmentary Figure S1

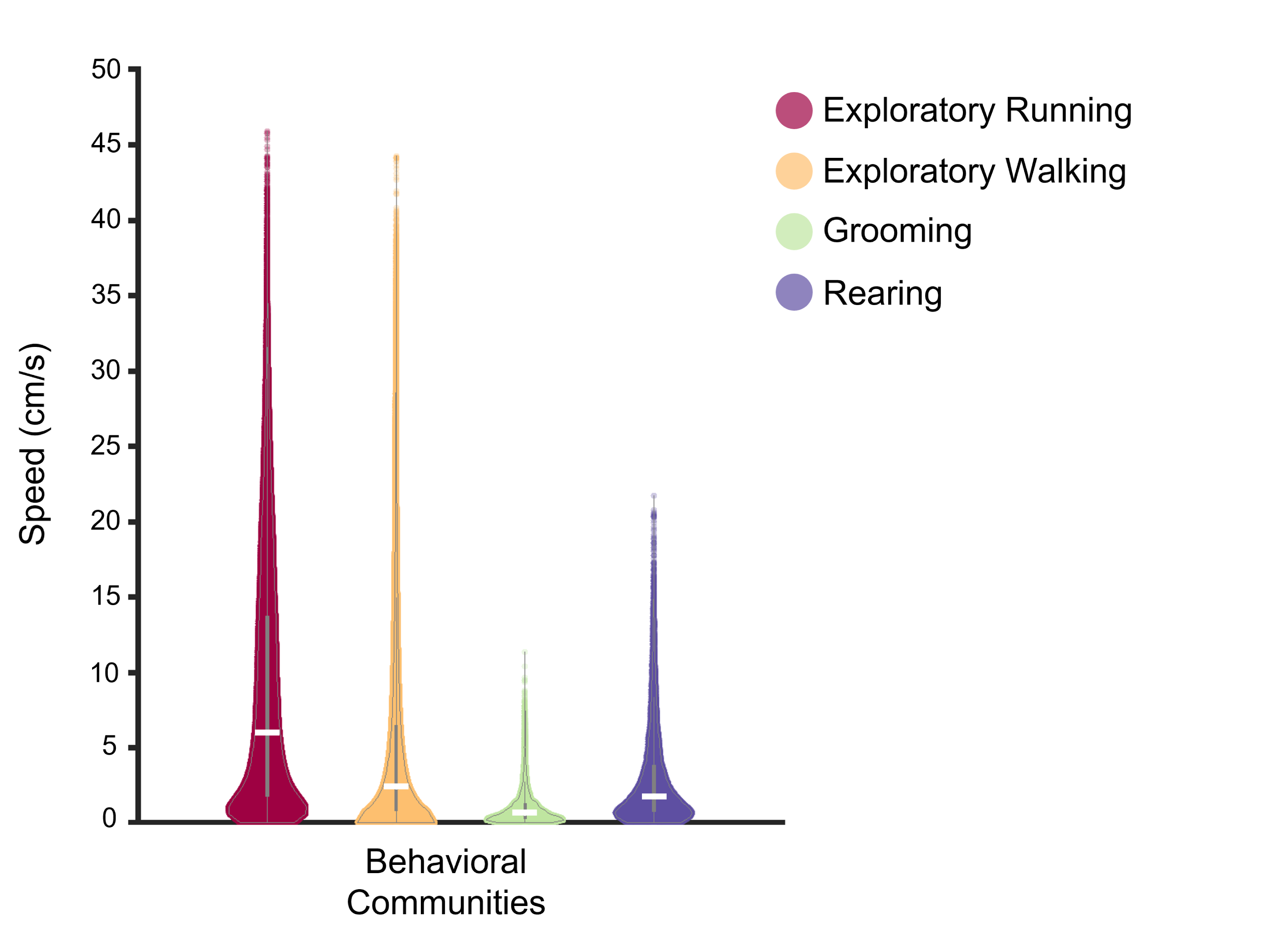
